# Methane-derived carbon flow through host-virus trophic networks in soil

**DOI:** 10.1101/2020.12.16.423115

**Authors:** Sungeun Lee, Ella T. Sieradzki, Alexa M. Nicolas, Robin L. Walker, Mary K. Firestone, Christina Hazard, Graeme W. Nicol

## Abstract

The concentration of atmospheric methane continues to increase with microbial communities controlling soil-atmosphere fluxes. While there is substantial knowledge of the diversity and function of organisms regulating methane production and consumption, the frequency and impact of interactions with viruses on their activity in soil is unknown. Metagenomic sequencing of soil microbial communities has enabled identification of linkages between viruses and hosts. However, determining host-virus linkages through sequencing does not determine whether a virus or a host are active. In this study, we identified active individual interactions *in situ* by following the transfer of assimilated carbon from active hosts to viruses. Using DNA stable-isotope probing combined with metagenomic analyses, we characterized methane-fueled microbial networks in acidic and neutral pH soils, specifically primary and secondary utilisers of carbon, together with the recent transfer of methane-derived carbon to viruses. Sixty-three percent of viral contigs from replicated soil incubations contained genes associated with known methanotrophic bacteria. Genomic sequences from ^13^C-enriched viruses were present in clustered regularly interspaced short palindromic repeats (CRISPR) arrays of multiple, closely-related *Methylocystis* populations, revealing differences in their history of viral interaction. Viruses infecting non-methanotrophic methylotrophs and heterotrophic predatory bacteria were also identified through the analysis of shared homologous genes, demonstrating that carbon is transferred to a diverse range of viruses associated with methane-fueled microbial food networks.

## Main text

Microorganisms play a central role in global carbon (C) biogeochemical cycling in soil systems. Soil is one of the most diverse habitats in the biosphere and can typically contain 10^9^ −10^10^ prokaryotic cells (Frossard et al., 2016) or viruses (Williamson et al., 2017) per g. Infection by viruses facilitates the horizontal transfer of genes and viral lysis acts as a control of host abundance and releases nutrients. In the marine environment, 20-40% of prokaryotes are lysed on a daily basis with the release of 150 Gt of carbon annually (Suttle, 2007). However, the role of viruses in influencing prokaryotic ecology in soil remains comparatively unknown (Emerson, 2019). In particular, difficulties remain in identifying the frequency of active interactions between native host and virus populations *in situ*, largely due to a lack of tools to study interactions within the highly complex and heterogeneous soil environment. While red-queen or ‘arms race’ dynamics have not yet been observed in natural soil populations as they have in marine systems (Ignacio-Espinoza et al., 2020), studies have shown viruses can coevolve with their hosts in soil and that hosts change in their susceptibility to infection (Gómez and Buckling, 2011). Shotgun sequencing of diverse soil microbial communities has enabled identification of linkages between viruses and hosts involved in carbon cycling both through identifying CRISPR spacer sequences in viral genomes and the presence of viral genes encoding enzymes involved in complex carbon degradation (Emerson et al., 2018). However, determining virus-host associations *in situ* with these methods does not elucidate the frequency of viral infections, with linkages potentially associated with populations not active under current conditions, or even relic DNA (Carini et al., 2017).

Methanotrophs are a critically important group in soil systems, removing 5% of atmospheric methane (Curry, 2007) and controlling fluxes to the atmosphere from methanogenic activity in anoxic compartments (Le Mer and Roger, 2001). Aerobic methanotrophs use CH_4_ for both carbon and energy requirements and key representatives in soil belong to the type I Gammaproteobacteria family *Methylococcaceae*, type II Alphaproteobacteria families *Methylocystaceae* and *Beijerinckiaceae*, and *Methylacidiphilaceae* of the Verrucomicrobia (Kneif, 2015). Soil pH is one of many factors influencing methanotroph activity, with type I and type II methanotrophs often dominating activity in neutral and acidic pH soils, respectively (Zhao et al., 2020). In addition, a wide variety of non-methanotrophic methylotrophs utilise methanol produced and excreted by methanotrophs, and together methanotrophic and other methylotrophic single carbon compound (C1)-utilising consortia assimilate methane-derived carbon in a variety of habitats (Chistoserdova et al., 2010).

A widely used technique for identifying active populations within a diverse microbial community in environmental samples, including methanotrophs, is DNA stable isotope probing (SIP) (Radajewski et al., 2002). Incorporation of a substrate with an enriched isotope can be traced in genomes of community members, demonstrating utilisation of a specific substrate linked to the associated functional process. As viruses are entirely composed of elements derived from a host cell, their production inside active hosts incorporating an isotopically-enriched substrate will also result in detectable viral isotopic enrichment (Pasulka et al., 2018). In this study we aimed to identify active virus-host interactions within a complex soil habitat by focussing on a taxonomically and functionally restricted group of organisms. By following ^13^C flow *in situ*, we aimed specifically to identify lytic DNA viruses of methanotrophs actively using CH_4_-derived C, including the identification of individual virus-host interactions, and potentially those actively infecting secondary utilisers such as non-methanotrophic methylotrophs.

After aerobically incubating pH 4.5 and 7.5 soils in the presence of ^12^C- or ^13^C-CH_4_, high buoyant density genomic DNA (>1.732 g ml^-1^) containing ^13^C-enriched or ^12^C-high GC mol% genomic DNA was recovered from triplicate incubations per isotope and soil via isopycnic centrifugation in CsCl gradients (Supplementary Fig. 1). Six metagenomes were produced from ^13^C isotopically-enriched DNA samples only (three pH 4.5, three pH 7.5; Supplementary Table 1). Concentrations of high buoyant density genomic DNA from ^12^C-CH_4_ incubations were too low for comparable shotgun sequencing. While this indicated minimal recovery of unenriched DNA in ^13^C-incubated samples, analysis of 16S rRNA gene amplicon libraries prepared from high buoyant density DNA of both ^12^C and ^13^C-CH_4_ incubations confirmed ^13^C-enrichment of C1-utilising populations (Supplementary Text, Supplementary Fig. 2).

Reads from individual metagenomes were assembled before taxonomic assignment of individual contigs. Reproducibly distinct communities were enriched in the two soils (Supplementary Fig. 2), with six bacterial families representing annotated contigs >5 kbp to which >1% of reads were mapped and all including known C1-utilising taxa (*Beijerinckiaceae, Bradyrhizobiaceae, Hyphomicrobiaceae, Methylococcaceae, Methylocystaceae* and *Methylophilaceae*). We resolved twenty-three medium and high-quality (Bowers et al., 2017) metagenome-assembled genomes (MAGs) (Supplementary Table 2), including 12 methanotrophs. Specifically, 3 MAGs represented gammaproteobacterial type I methanotrophs (*Methylobacter*) and 9 MAGs represented alphaproteobacterial type II methanotrophs (*Methylocystis, Methylosinus* or *Methylocapsa*). Secondary utilisers of methane-derived organic carbon were also identified with 9 MAGs associated with established or putative non-methanotrophic methylotrophs, lacking methane oxidation machinery but capable of utilising methanotroph-derived methanol (see Supplementary Text). These included representatives of the *Gemmatimonadales, Hyphomicrobium, Herminiimonas* and *Rudaea*, the latter two, to our knowledge, not having been previously associated with methylotrophy but possessed predicted methanol and formate dehydrogenases (Supplementary Table 2). Two MAGs represented strains of *Bdellovibrio* and *Myxococcus*, known predatory bacteria, indicating that growing methylotrophic populations were preyed upon (Pérez et al., 2018).

Lytic virus populations linked to C1-hosts were analysed using metagenome viral contigs (mVCs), predicted using established tools. Using contigs >10 kbp (Roux et al., 2019) VirSorter (Roux et al., 2015) predicted 270 metagenome viral contigs (mVCs) with a further 4 ‘likely’ mVCs predicted uniquely by DeepVirFinder (Ren et al., 2020) (see Supplementary Text), together representing 227 viral operational taxonomic units (vOTUs) (Paez-Espino et al., 2017). Analysis of the normalised read mapping for mVCs demonstrated that, as with the bacterial communities, active ^13^C-enriched viral populations were reproducibly distinct between acidic and neutral pH soils (Supplementary Fig. 3).

mVCs were linked to host bacteria using three different approaches: identifying incorporation of viral DNA into spacers of bacterial CRISPR arrays, similarity of homologous genes possessed by both host and virus contigs, and *k*-mer similarity between potential host and virus contigs, the last approach being considered only partially successful (see Supplementary Text). CRISPR arrays were identified in 3 of 23 MAGs, each associated with the genus *Methylocystis* or *Methylosinus* of the *Methylocystaceae* (Fig. 1). In the acidic soil, complete CRISPR arrays of growing methanotrophs were associated with two *Methylocystis* MAGs (MAG identifiers 5 and 6) sharing 79.2% average nucleotide identity (ANI) and likely representing different species (Jain et al., 2018). A further six CRISPR arrays were identified in unbinned bacterial contigs all possessing the same direct repeat (DR) sequence. These eight arrays varied in size ranging from 9 to 114 DRs and contained a total of 432 spacers, and were in the same size range of *Methylocystaceae* CRISPR arrays from sequenced genomes (Supplementary Table 3). Comparison of spacer incorporation between arrays revealed that these multiple closely-related populations had different histories of viral interaction and subsequent spacer incorporation. Genome sequences from ^13^C-enriched viral populations were represented by seven mVCs and matched 29.5% of spacers. In addition, 7.9% of spacers possessed a one nucleotide mismatch, all of which represented a synonymous substitution, indicating that variation was the result of mutations in viral genomes increasing their ability to evade CRISPR-CAS defense systems or genetic variation in closely related viral populations. Only three pairs of spacers were identical, with each pair member located on a different array. Variation in virus host range was also observed, with 3 and 2 mVCs linked to only one or both *Methylocystis* MAGs, respectively.

**Fig. 1.**
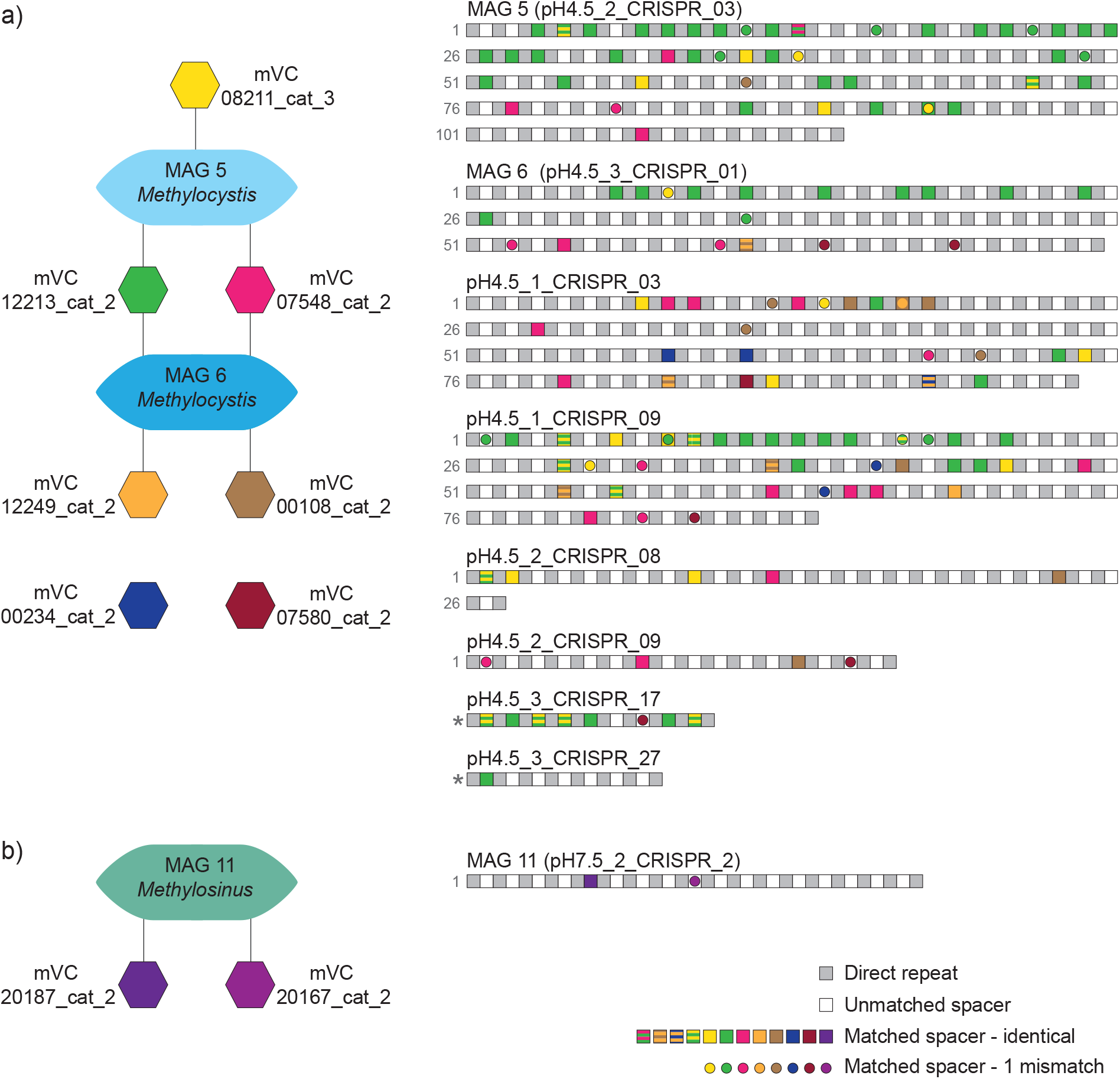
Linkage of active ^13^C-enriched viruses to *Methylocystaceae* populations in soil by comparison of spacer sequences in CRISPR arrays. a) Distribution of spacers from 8 mVCs in *Methylocystis* CRISPR arrays (MAGs 5, 6 and six unbinned contigs). b) CRISPR array of *Methylosinus* MAG 11 containing spacers linked to two mVCs. CRISPR array names describe the individual soil microcosm that the contig was recovered from. DRs for complete arrays are numbered (in grey), with the spacer after DR 1 being the most recently incorporated. Two partial arrays are denoted with an *. Spacers with 100% identity or 1 mismatch to sequences in mVCs are represented by colour-coded squares and circles, respectively, with stripes representing sequences found in two different mVCs.

Surprisingly, a large number of spacers in individual *Methylocystis* CRISPR arrays were linked to the same virus, with up to 31 being homologous to protospacer sequences in one mVC. To provide support that these multiple spacers were derived from *Methylocystis*-associated viruses, mVCs were examined for host-specific conserved protospacer-adjacent motif (PAM) sequences (Mojica et al., 2009). Consistent with the identification of genuine protospacers, 138 of linked 146 spacers (i.e. all possessing ≤ 1 mismatch) had the conserved PAM sequence ‘TTC’ (target-centric orientation) (Leenay and Beisel, 2017). The variation in spacer position between the arrays revealed temporal differences in virus infections. For example, the most recently integrated spacer in 3 of 8 different *Methylocystis* CRISPRs was derived from a ^13^C-enriched virus represented by mVC_12213_cat2, suggesting the possibility of incorporation occurring during the incubation of the experiment.

Further analysis of all CRISPR arrays (i.e. including those in unbinned contigs) linked to ^13^C-enriched viruses revealed that the majority of viruses were associated with methanotrophic populations (Fig. 2a). In total, 11 different variants were identified (i.e. each having a unique DR sequence) with 9 linked to *Methylocystaceae* or *Methylococcaceae* populations. DR sequences generally possessed high sequence similarity to those in CRISPR arrays from cultivated strain genomes of the same family, although only CRISPR array 6 had a DR sequence that was identical (Supplementary Table 3). Individual DR variants were restricted to either pH 4.5 or 7.5 soil. Using 100% sequence identity in searches between CRISPR spacer and mVC protospacer sequences, 19 VirSorter-predicted mVCs were linked to all CRISPR array variants. In addition, analysis of shorter mVCs ranging 5-10 kbp identified two additional linked mVCs (mVC_08964_cat.3 (9.8 kbp) and mVC_28139_DVF (5.1 kbp)). One third of CRISPR linked-mVCs were categorized at the lowest level of confidence (i.e. category-3 by VirSorter (Roux et al., 2015) or ‘possible’ by DeepVirFinder (Ren et al., 2020), suggesting that retaining only higher confidence contigs may exclude a substantial proportion of *bona fide* methanotroph virus-derived contigs in uncharacterised environments such as soil.

**Fig. 2.**
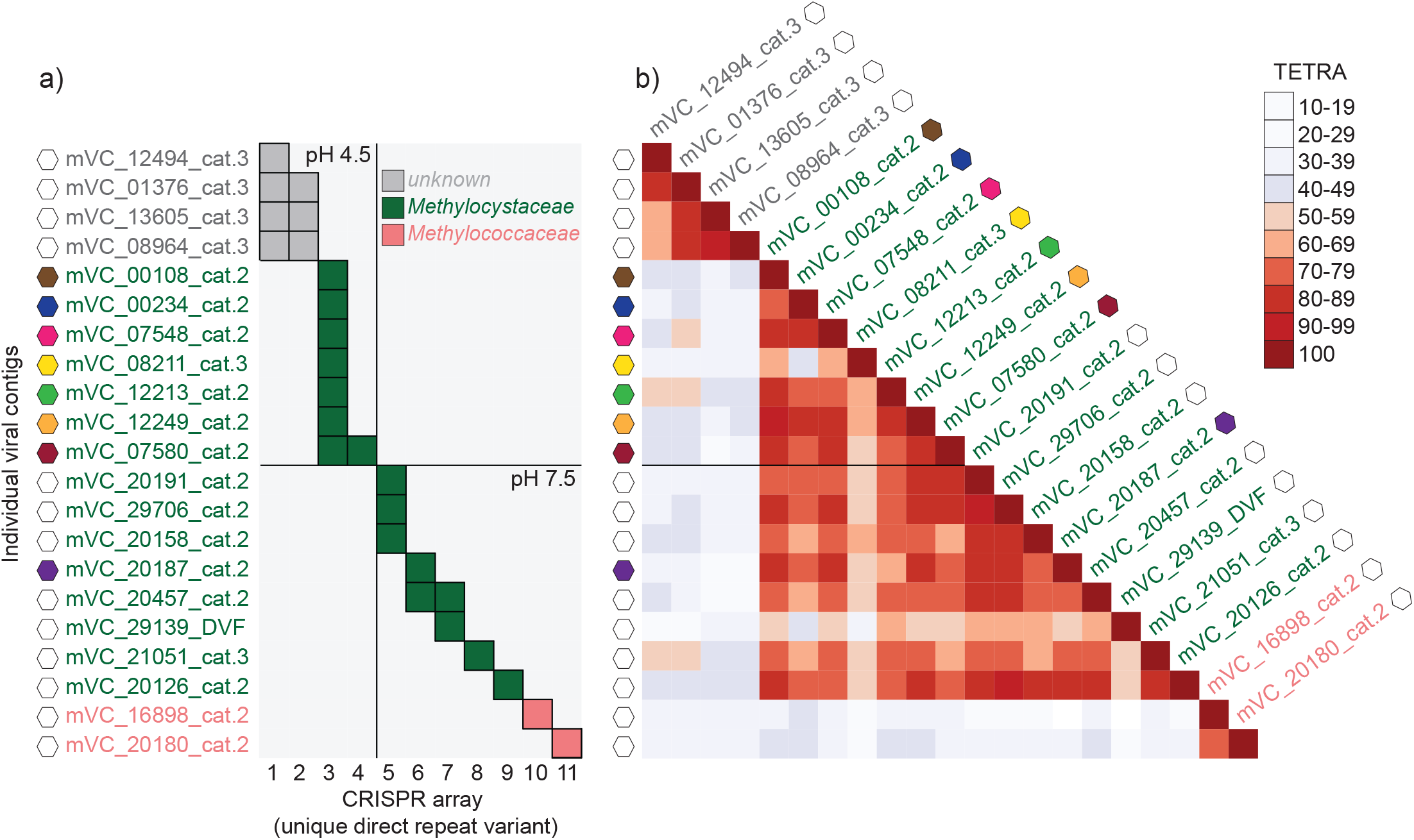
Linkages of ^13^C-enriched viruses to CRISPR arrays in pH 4.5 and 7.5 soil. a) Presence of spacers from 21 mVCs in 11 different CRISPR array variants (unique DR sequence). Taxonomic affiliation of CRISPR arrays to host families was determined by phylogenomic analysis of affiliated MAGs (3, 6) or unbinned contigs (2-5, 7-9), or inferred from shared homologues between linked mVCs and bacterial genomes (10, 11). All mVCs were >10 kbp except mVC_08964_cat.3 (9.8 kbp) and mVC_28139_DVF (5.1 kbp). These two mVCs were also the only two predicted using DeepVirFinder, with calculated probabilities describing ‘likely’ and ‘probable’ viruses, respectively. b) TETRA correlation coefficients between 21 CRISPR-linked mVCs. Colour-coded hexagon symbols denote linkage to MAG-associated CRISPR arrays as per Fig. 1.

Analysis of tetranucleotide frequencies (TETRA) (Wang et al., 2017) clustered the 21 mVCs into three groups that were associated with the *Methylocystaceae, Methylococcoceae* and an unknown group (Fig. 2b). The majority of viruses infected members of the *Methylocystaceae* family, with those infecting populations of the *Methylocystis* and *Methylosinus* genera restricted to acidic and neutral pH soils, respectively. TETRA correlation coefficients of all *Methylocystaceae*-linked viruses were in the same range both within and between either genus, suggesting co-evolution with their host rather than genetic drift and divergence was the primary mechanism for defining specific associations with *Methylocystis* or *Methylosinus* strains.

Identification of homologous genes in mVCs that were shared with prokaryotic genomes were always consistent with host-virus linkages established using spacer sequences from MAG CRISPR arrays. Specifically, BLASTp searches of genes present in the 9 mVCs linked to *Methylocystaceae* MAGs via CRISPR spacer sequences all contained ‘best hits’ (identity >30%, e-value <10^−5^, bit score >50 and query cover >70%) to a minimum of 5 homologues also found in *Methylocystaceae* genomes. This was therefore used as a criterion for establishing host-virus linkages. Sixty-three percent of mVCs contained a homologue that was linked to genomes of known C1-utilising bacteria, with 35% linked specifically to populations from the *Methylocystaceae, Methylococcaceae* or *Hyphomicrobiaceae* (Fig. 3a). While analysis of bacterial homologues in mVCs identified the taxonomic family of the assumed dominant host, they also indicated that individual viruses may infect hosts of other families of the same taxonomic order, including those at other trophic levels. Specifically, within the *Rhizobiales*, mVCs linked to *Methylocystaceae* also contained homologues shared with *Bradyrhizobiaceae, Methylobacteriaceae* and *Rhizobiaceae* (Fig. 3b), indicating that viruses of methanotrophs may also infect non-methanotrophic methylotrophs that are active at the same time.

**Fig. 3.**
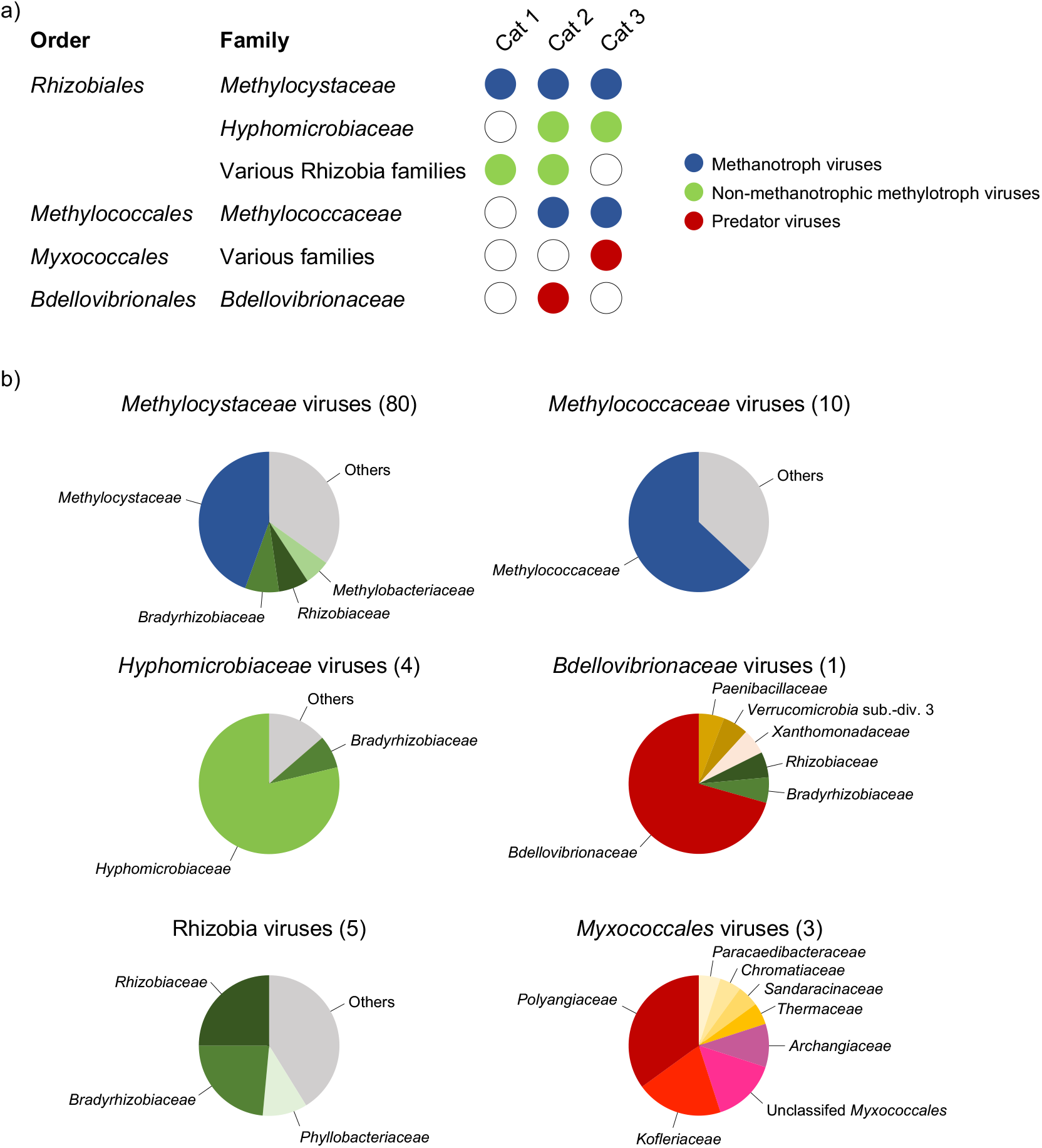
Linkage of ^13^C-enriched viruses to methanotrophic, methylotrophic and predator bacterial host populations through identification of shared homologous genes. a) Association of viruses with different bacterial families and functional groups inferred from the presence of ≥5 shared homologous genes in category-1, −2 and −3 VirSorter-predicted mVCs. b) Proportion of homologues in methanotroph, non-methanotrophic methylotroph or predator viruses linked to individual bacterial families. Each chart summarises those mVCs that all contain ≥5 homologues to one family (number of mVCs given in parentheses) but with other taxonomic linkages also given. ‘Other’ describes the proportion found in families each represented by less than <5% of homologues or those not annotated to the family level.

CH_4_-derived C was also transferred to viruses of secondary or tertiary utilisers. One group of mVCs were linked to methylotrophic *Hyphomicrobiaceae* and a second to a phylogenetically diverse range of nitrogen-fixing Rhizobia i.e. *Bradyrhizobiaceae, Phyllobacteriaceae* and *Rhizobiaceae*. These lineages contain known methylotrophs, methanol dehydrogenases have been identified in a range of rhizobial species (Huang et al., 2019) and these mVCs also contained homologues found in the genomes of nodulating *Methylobacterium* strains (Green and Ardley, 2018). Viruses of predatory *Bdellovibrio* and *Myxococcales* bacteria were predicted, consistent with the recovery of corresponding bacterial MAGs in ^13^C-enriched DNA. One mVC (20210-cat_2) was linked to the genus *Bdellovibrio* (sharing 11 of 67 mVC genes) and three category-3 mVCs (i.e. possible viruses) was linked to *Myxococcales* populations, containing gene homologous to four families within the order. The high isotopic labelling of both heterotrophic predators (with no identifiable C1-utilising capability) and their viruses indicate that the predators were feeding primarily on C1-utilisers, as carbon incorporated by feeding on unlabelled bacteria would dilute the enrichment in predators. As such, it also indicates that predatory bacteria may have preference for preying upon growing populations rather than the non-C1-utilising majority.

Gene-sharing network analysis of mVCs with viruses in the NCBI RefSeq database and other metagenome studies were analysed using vConTACT 2.0 (Jang et al., 2019). Any linkages with RefSeq viruses typically had low scores (i.e. sharing a low number of homologues) and were linked to viruses of hosts that were inconsistent with our homologue-based predictions (Supplementary Table 4). No linkages were observed with recently reported giant viruses of methanotrophs in freshwater lakes (Chen et al., 2020). However, in a recent study of 197 metagenomes from Swedish peatland soil, Emerson et al. (2018) identified 13 viruses linked to methanotrophs. Intriguingly, 8 of these were linked in our viral gene-sharing network, with both studies predicting *Methylocystaceae* hosts using different methods of host annotation (Supplementary Text, Supplementary Fig. 5) and revealing the distribution of specific *Methylocystaceae*-associated viral groups present in different geographical areas and soil types. Analysis of gene-sharing networks of mVCs from this study indicated that there were two distinct *Methylocystaceae*-linked viral clusters which also varied in their distribution in both soils. Specifically, one cluster was associated with low pH only whereas the second cluster contained viruses found in both pH 4.5 and 7.5 soils, including those linked by CRISPR spacer sequences. Individual networks of *Methylococcaceae-* and rhizobia-associated mVCs were also identified, typically associated with one of the two soils of contrasting pH. Taxonomically-linked mVCs with ≥5 homologues were consistently placed in networks with other mVCs containing 1-4 homologues from the same methylotrophic families, confirming host linkage to a larger number of mVCs.

mVCs contained 8,174 genes, with 49.6% (4,054) annotated representing 606 unique functions. Of these, genes encoding viral proteins accounted for 9.8% (397 genes) and included major capsid proteins, tail proteins, integrases, portal proteins and terminases. Bacterial proteins used for viral replication accounted for 5.1% (206 genes). A number of metagenomic studies have demonstrated that viruses can possess genes encoding sub-unit C of ammonia or particulate methane monooxygenases as auxiliary metabolic genes (AMGs) (Chen et al., 2020; Ahlgren et al., 2019) which are also typically found as isolated genes in genomes in addition to being present in clusters or operons encoding A and B sub-units (Nicol and Schleper, 2006). In this study, one low confidence mVC (7.3 kbp, category-3) was identified as containing an isolated *pmoC* gene that was phylogenetically related to growing *Methylocystis* populations but was distinct from *pmoC* sequences found in viruses associated with freshwater *Methylocystis* populations (Chen et al., 2020) (Supplementary Fig. 6).

In summary, these results demonstrate that by following carbon flow, viruses and hosts associated with a critical biogeochemical process can be identified at the scale of individual populations, and currently active interactions at different trophic levels examined within the highly complex soil environment. Type I and II methanotrophs interact with evolutionarily distinct groups of viruses and the composition of CRISPR arrays of *Methylocystaceae* reveal that they have a continual dynamic interaction with individual viruses. Analysis of shared homologues in individual viral genomes show that they may interact with host populations at different trophic levels within a methane-fuelled network.

## Methods

### Soil microcosms

Triplicate soil samples were collected in February 2018 at 1 m intervals from the upper 10 cm of pH 4.5 and 7.5 soil sub-plots of a pH gradient maintained since 1961 and under an 8-year crop rotation (SRUC, Craibstone Estate, Aberdeen, Scotland; UK grid reference NJ872104) (Kemp et al., 1992). The crop at the time of sampling was potatoes. Soil (podzol, sandy-loam texture) was sieved (2 mm mesh size) and microcosms established in triplicate for each soil pH and isotope in 144 ml serum bottles containing 14.30 g soil (10 g dry weight equivalent) with a 30% volumetric water content, equivalent to ∼60% water-filled pore space. Bottles were capped and established with a 10% (v/v) ^12^C-CH_4_ or ^13^C-CH_4_ (Sigma-Aldrich) headspace (99% atom enriched), re-opening every 10 days to maintain aerobic conditions before sealing and re-establishing CH_4_ headspace concentrations. Microcosms were incubated at 25°C and destructively sampled after 30 days with soil archived immediately at −20°C.

### DNA-SIP

Genomic DNA was extracted from 0.5 g soil samples using a CTAB buffer phenol:chloroform: isoamyl alcohol bead-beating protocol and subjected to isopycnic centrifugation in CsCl gradients, recovery and purification as previously described (Nicol and Prosser, 2011). Briefly, 6 ug of genomic DNA was added to 8 ml CsCl-Tris EDTA solution (refractive index (RI) of 1.4010; buoyant density of 1.71 g ml^−1^) in polyallomer tubes before sealing and ultracentrifugation at 152,000 × *g* (50,000 rpm) in a MLN80 rotor (Beckman-Coulter) for 72 h at 25°C. CsCl gradients were fractionated into 350 ul aliquots using an in-house semi-automated fraction recovery system before determining RI and recovering DNA. The relative abundance of bacterial 16S rRNA and methanotrophic *pmoA* genes in genomic DNA distributed across the CsCl gradients was determined by qPCR in a Corbett Rotor-Gene 6000 thermocycler (Qiagen) using primer sets P1(341f)/P2(534r) (Muyzer et al., 1993) and A189F/A682R (Holmes et al., 1995) respectively. Twenty-five μl reactions contained 12.5 μl 2X QuantiFast SYBR Green Mix (Qiagen), 1 μM of each primer, 100 ng of T4 gene protein 32 (Thermo Fisher), 2 μl of standard (10^8^-10^2^ copies of an amplicon-derived standard) or 1/10 diluted DNA. Thermocycling conditions consisted of an initial denaturation step of 15 min at 95°C for both assays followed by 30 cycles of 15 s at 94°C, 30 s at 60°C, 30 s at 72°C for the 16S rRNA gene assay or 60 s at 94°C, 60 s at 56°C, 60 s at 72°C for the *pmoA* gene assay, followed by melt-curve analysis. All assays had an efficiency between 93-97% with an r^2^ value >0.99. Genomic DNA from four fractions with a buoyant density >1.732 g ml^-1^ were then pooled for each ^12^C- and ^13^C-CH_4_-incubated replicate for 16S rRNA gene amplicon sequence and metagenomic analysis.

### Metagenome sequencing, assembly & annotation

Library preparation and sequencing was performed at the Joint genome Institute (JGI), Berkeley, USA. Libraries were produced from fragmented DNA using KAPA Biosystems Library Preparation Kits (Roche) and quantified using KAPA Biosystems NGS library qPCR kits. Indexed samples were sequenced (2 x 150 bp) on the Illumina NovaSeq platform with NovaSeq XP v1 reagent kits and a S4 flowcell. Raw reads were processed with JGI’s RQCFilter2 pipeline that utilised BBTools v38.51 (Bushnell, 2016). Reads containing adapter sequences were trimmed and those with ≥3 N bases or ≤ 51 bp or ≤ 33% of full-read length were removed along with PhiX sequences using BBDuk, and reads mapped to human, cat, dog or mouse references at 95% identity were removed using BBMap. *De novo* contig assembly of the 100 - 196 million quality-controlled reads per metagenome was performed using MetaSPAdes v3.13.0 (Nurk et al., 2016). The 1 - 2 million contigs per metagenome were then concatenated together, and contigs larger than 5 kbp were dereplicated at 99% average nucleotide identity (ANI) using PSI-CD-HIT v4.6.1 (Fu et al., 2012) and binned using MetaWRAP v1.2.1 (Uritskiy et al., 2018) (Supplementary Table 5). Bin completion and contamination was determined by CheckM v1.0.12 (Parks et al., 2015). Taxonomic annotation of contigs was performed using Kaiju (Menzel et al., 2016) with the NCBI RefSeq database (Release 94; 25 June 2019) (O’Leary et al., 2016) and MAGs using GTDB-Tk v0.3.2 (Chaumeil et al., 2019) with the Genome Taxonomy Database (release 89, 21 June 2019) (Parks et al., 2018). Protein sequence annotation was performed using InterProScan 5 (e-value <10^−5^) (Jones et al., 2015). Pairwise ANI comparison of MAGs was calculated using FastANI (Jain et al., 2019).

### Amplicon sequencing and analysis

16S rRNA genes were amplified using primers 515F/806R (Walters et al., 2015) followed by library preparation and sequencing on an Illumina MiSeq sequencer as previously described (Finn et al., 2020). Reads with a quality score <20 and length < 100 bp were discarded using FASTX-Toolkit v0.0.13 (http://hannonlab.cshl.edu/fastx_toolkit/). High-quality reads were merged using PANDAseq v2.11 (Masella et al., 2012), and denoising and chimera removal performed with UNOISE3 (Edgar, 2016). Amplicon sequence variants (ASVs) were annotated using the RDP classifier v2.11 (Wang et al, 2007). Non-metric multidimensional scaling of Bray-Curtis dissimilarity derived from the relative abundance of ASVs was performed with the metaMDS function in the vegan package (Oksanen et al., 2019) in R v3.6.0.

### Virus prediction

Metagenomic viral contigs (mVCs) were predicted from 9,190 contigs >10 kbp using VirSorter (Roux et al., 2015), retaining non-prophage category-1, −2 or −3 mVCs, representing “most confident”, “likely” and “possible”. DeepVirFinder (Ren et al., 2020) was also used to predict mVCs from contigs >10 kbp, with those with a *p*-value <0.05 and a score ≥0.9 or ≥0.7, representing “confident” and “possible”, respectively (Supplementary Table 6). The relative abundance of each mVC in the six metagenomes was determined using the MetaWRAP-Quant_bins module (Uritskiy et al., 2018) and a heatmap produced using the heatmaply package in R v3.6.0.

### Virus-host linkage

CRISPR arrays within MAGs and unbinned contigs were identified using the CRISPR Recognition Tool v1.2 (Bland et al., 2007) (Supplementary Table 7). DR and spacer sequences were extracted before performing 100% identity searches against positive and negative strands to identify MAGs or contigs with direct repeats and the viral origin of spacers using Seqkit commands (Shen et al., 2016). After identification of matched spacer sequences in mVCs, 10 nucleotides before and after the spacer sequence were extracted to identify associated host-specific PAM sequences. Conserved and variant PAM sequences were manually identified. Correlation coefficients of pairwise comparison of the tetra-nucleotide frequencies (TETRA) between unique CRISPR-associated mVCs were calculated using Python package pyani v0.2.10 (Pritchard et al., 2016). To identify homologous genes shared between CRISPR-linked viruses and hosts, gene prediction was performed using Prodigal v2.6.3 (Hyatt et al., 2010) with the -p meta option followed by protein alignment with Blastp (identity >30%, e-value < 10^−5^, bit score >50 and query cover >70%) and protein sequence annotation using InterProScan 5 (e-value <10^−5^). Gene homology between all mVCs and prokaryotes in the NCBI nr database was determined using Diamond Blastp (e-value <10^−5^) (Buchfink et al., 2015). Virus-host prediction using k-mer frequencies was performed with WIsH v1.0 (p-values <0.05) (Galiez et al., 2017). Networks based on shared gene content was constructed using vConTACT 2.0 (Jang et al., 2019) with the NCBI RefSeq database (Release 94; 25 June 2019).

### Phylogenetic analysis of PmoC and PxmC protein sequences

Maximum likelihood analysis of inferred protein sequences of membrane-bound monooxygenase C sub-units from methanotroph MAGs and reference sequences (Supplementary Table 8) was performed on unambiguously aligned sequences using PhyML 3.0 (Guindon et al., 2010) with automatic model selection (LG substitution, gamma distribution (0.06) and proportion of invariable sites (0.087) estimated). Bootstrap support was calculated from 100 replicates.

## Data availability

Metagenome sequence reads are deposited under NCBI BioProject accession numbers PRJNA621430 - PRJNA621435. Metagenome draft assemblies are accessible through the JGI Genome Portal (DOI: 10.25585/1487501). Amplicon sequence data is deposited in the NCBI Sequence Read Archive with BioProject accession number PRJNA676099.

## Acknowledgments

The sequencing data were generated under JGI Community Science Program proposal 503702 awarded to GWN and CH. The work conducted by the U.S. Department of Energy Joint Genome Institute, a DOE Office of Science User Facility, is supported by the Office of Science of the U.S. Department of Energy under Contract No. DE-AC02-05CH11231. This work was funded by an AXA Research Chair awarded to GWN and a France-Berkeley Fund grant (2018-2019) awarded to GWN and MKF. The pH gradient experiment is funded through the Scottish Government RESAS 2016-2021 programme. The authors would like to thank Dr. Joanne Emerson for valuable discussion.

## Author contributions

The research program was conceived by and funded from grants awarded to GWN, CH and MF. SL, CH and GWN designed the experiment and wrote the manuscript. SL performed experiments and analyses. ES, AN and MF advised on bioinformatic approaches, discussed data and commented on the manuscript. RW coordinated soil sampling and commented on the manuscript. All authors approved the manuscript.

---

**Supplementary Fig. 1.**
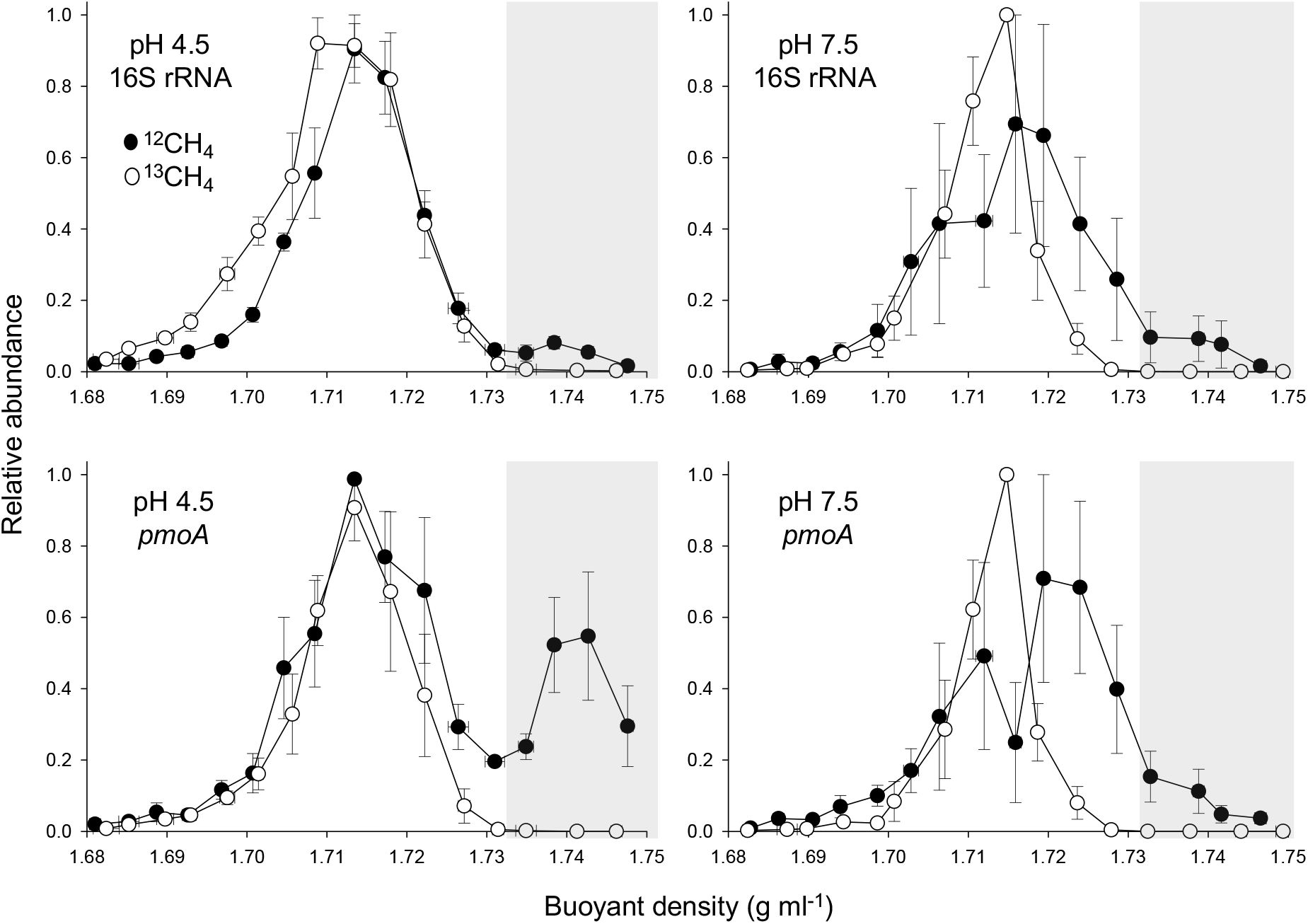
Buoyant density distribution of genomic DNA from total bacterial 16S rRNA genes and methanotroph communities possessing particulate methane monooxygenase sub-unit A (*pmoA*) genes after isopycnic centrifugation in CsCl gradients. Genomic DNA was extracted from triplicate pH 4.5 and 7.5 soil microcosms incubated with a 10% ^12^C- or ^13^C-CH_4_ headspace. Vertical error bars are the standard error of the mean relative abundance and horizontal bars (mostly smaller than the symbol size) the standard error of the mean buoyant density of individual fractions from three independent CsCl gradients, each representing an individual microcosm. The four fractions with the highest buoyant density (highlighted by grey area) were pooled for each replicate microcosm.

**Supplementary Fig. 2.**
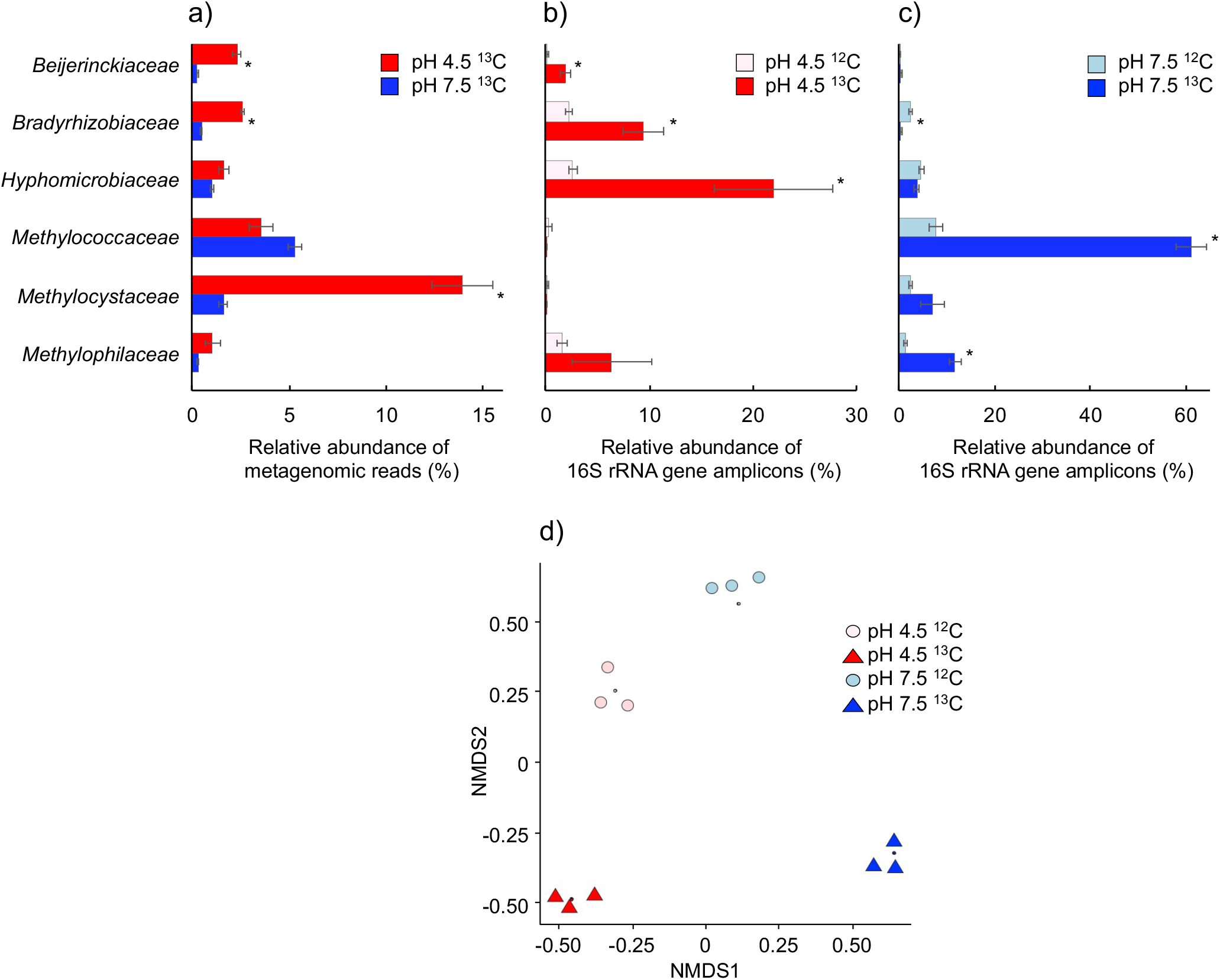
Taxonomic affiliation of metagenome reads and 16S rRNA gene amplified sequence variants (ASVs) derived from high buoyant density DNA from triplicate pH 4.5 and pH 7.5 soil microcosms after incubation with ^12^C- or ^13^C-CH_4_. a) Relative abundance of metagenome sequences mapped to contigs of families that recruited ≥1% reads in either soil. Reads were mapped to annotated contigs ≥5 kbp from ^13^C-incubated microcosms only. b) and c) Relative abundance of 16S rRNA gene ASVs derived from the six dominant families in ^12^C- or ^13^C-CH_4_ incubations of pH 4.5 and 7.5 soil, respectively. d) Non-metric multidimensional scaling of Bray-Curtis dissimilarities derived from the relative abundance of annotated 16S rRNA gene ASVs. Due to the close overlap of replicates (small symbols), samples were resolved for visualisation using a jitter function (large symbols). Significant differences between samples are indicated with * (*p* <0.05, two-sample Student’s t-test or Welsch’s t-test when variances were not homogenous).

**Supplementary Fig. 3.**
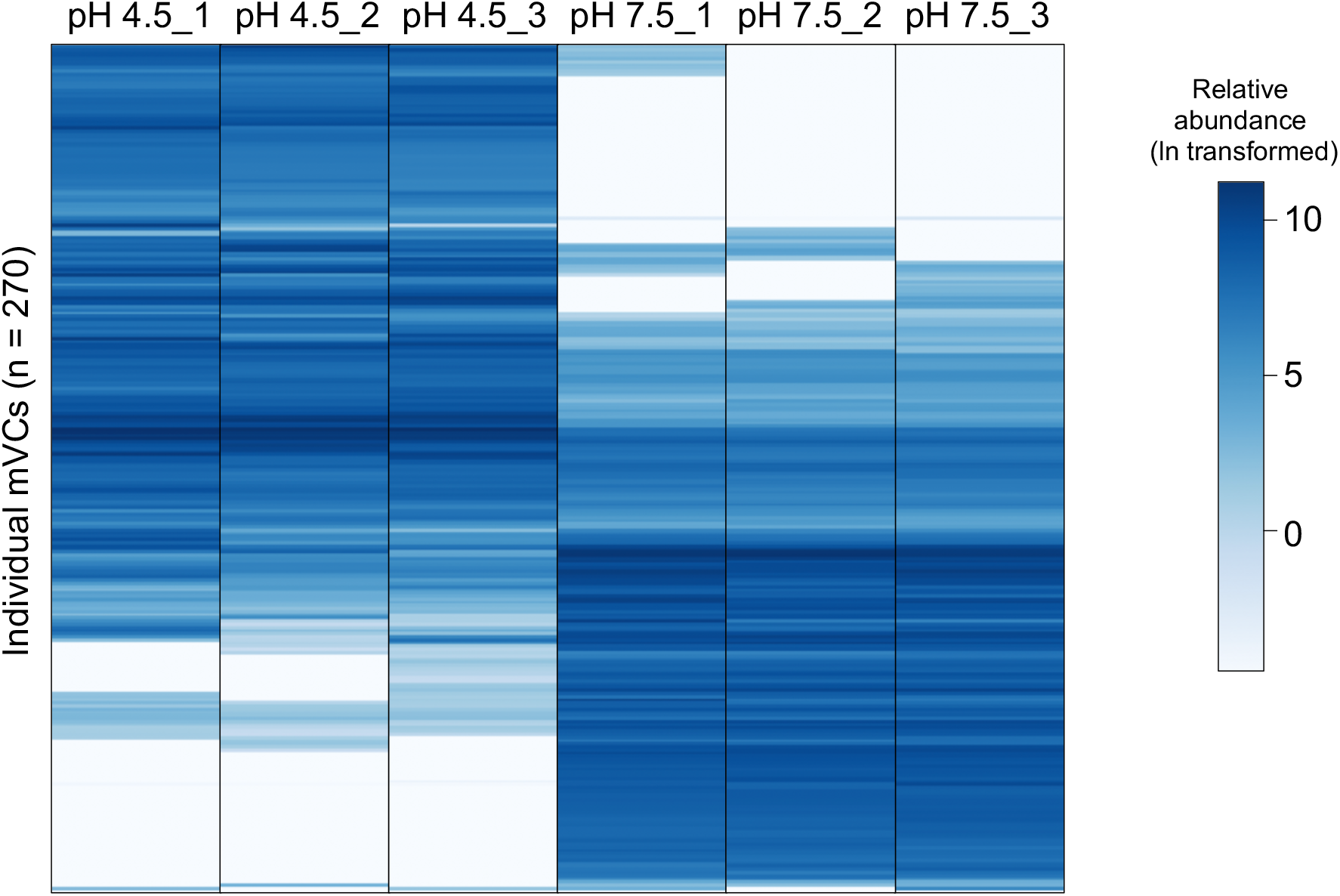
Heat-map displaying the relative abundance of 270 mVCs >10 kbp in length from ^13^C-enriched viral DNA derived from triplicate pH 4.5 and pH 7.5 soil microcosms. The values of normalised relative abundance are presented as reads per kbp after ln transformation.

**Supplementary Fig. 4.**
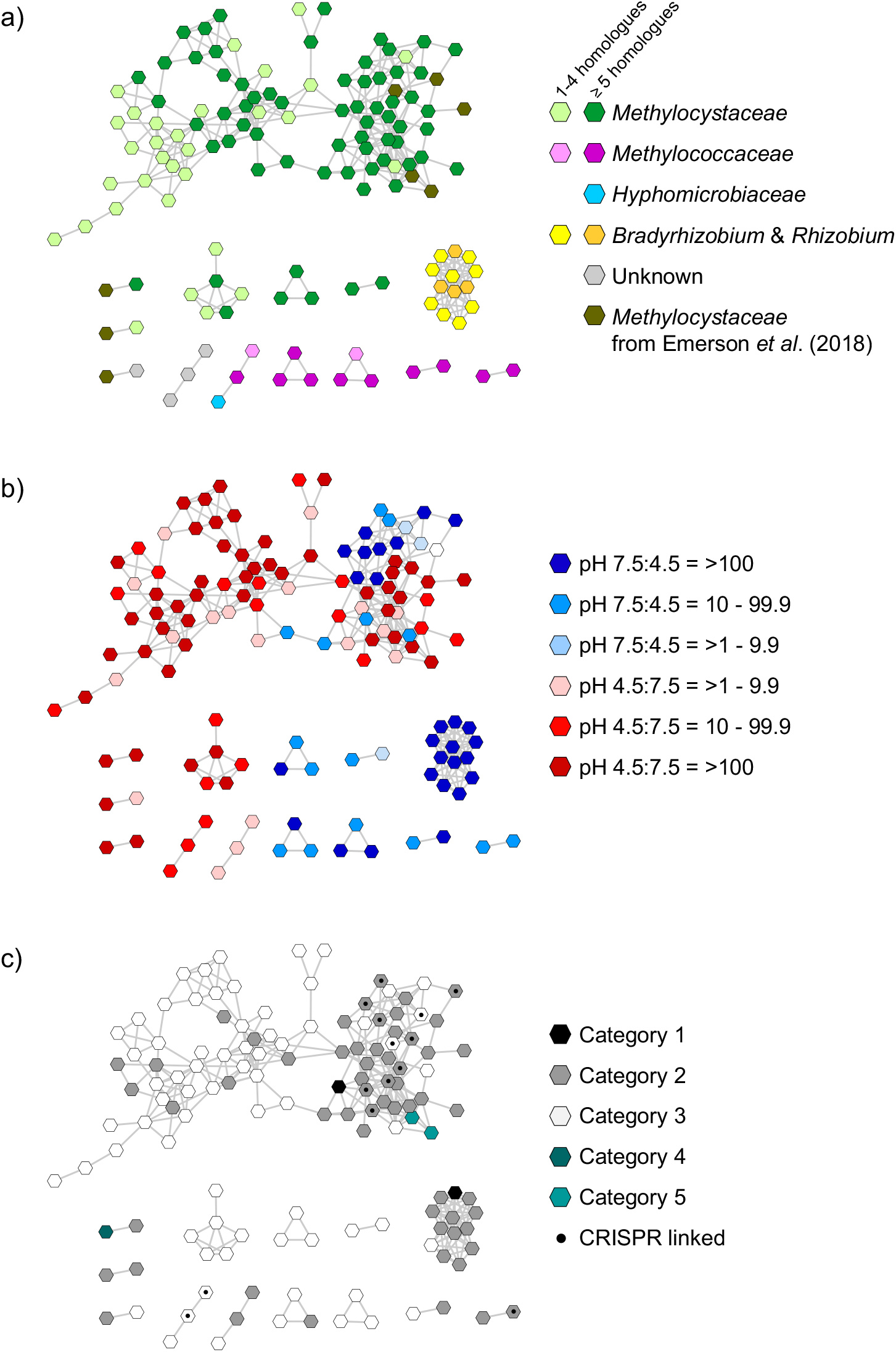
Gene sharing network analysis of mVCs (one representative per vOTU) from ^13^C-enriched viruses in pH 4.5 and 7.5 soils. a) Taxonomic affiliation of hosts predicted by homologue analysis, with mVCs containing ≥5 or <5 linked homologues highlighted. Eight mVCs from peatland soils linked to the *Methylocystaceae* (Emerson *et al*., 2018) are also shown. b) Distribution of mVCs in pH 4.5 and 7.5 soil determined by the mean ratio of normalised relative abundance from triplicate samples. Seven mVCs from peatland were only found in soils with pH ≤ 4.7. c) VirSorter category prediction and linkage to CRISPR arrays via spacer sequence analysis.

**Supplementary Fig. 5.**
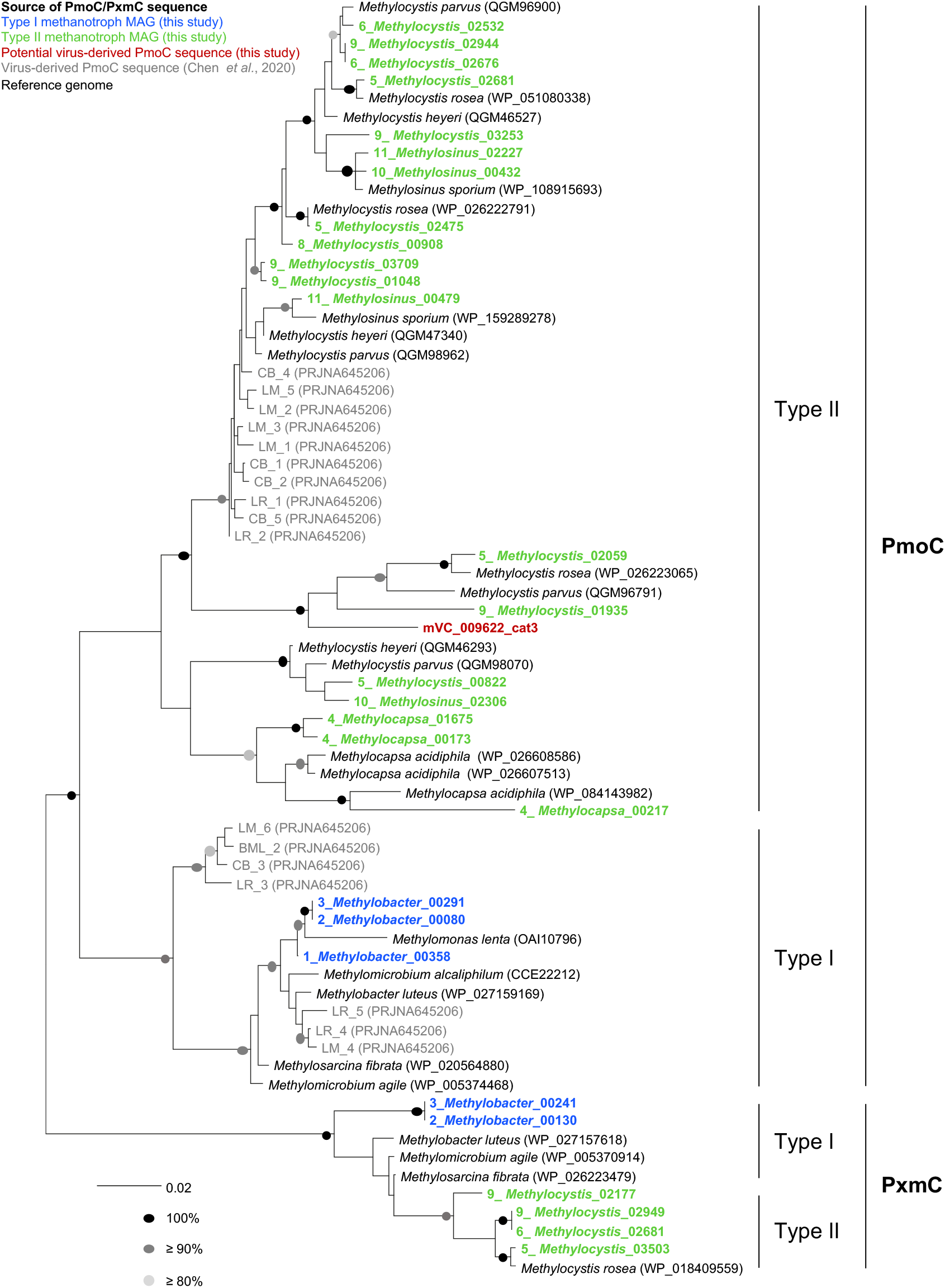
Maximum likelihood phylogenetic tree of derived amino acid sequences of PmoC and PxmC found in three type I and seven type II methanotroph MAGs and one potential viral-derived contig. MAG-derived sequences are described by MAG number, genus and contig identifier (see Supplementary Table 8). Sequences in reference methanotroph genomes and freshwater-derived viruses (Chen et al., 2020) were included with NCBI accession numbers given in parenthesis. Circles at nodes describe percentage bootstrap support from 100 replicates and the scale bar represents 0.02 changes per amino acid position.

## Supplementary text

### Comparison of 16S rRNA gene amplicon and metagenomic libraries of high buoyant density DNA from ^12^C and ^13^C-CH_4_ incubations

The relative abundance of annotated contigs belonging to different families in replicate metagenomic libraries was reproducible and distinct between acidic and neutral pH soils (Supplementary Figure 2a). Twenty of the 23 medium and high-quality MAGs recovered in ^13^C-derived metagenomic libraries were from methanotrophic or non-methanotrophic methylotrophic populations and therefore consistent with targeting a methane-fuelled community using stable isotope probing (Supplementary Table 2). Nevertheless, comparison with equivalent ^12^C incubations was performed as per standard practice with DNA-SIP experiments^1^. Genomic DNA was recovered and purified from high buoyant density fractions (>1.732 g ml^-1^) from triplicate microcosms of both ^12^C- and ^13^C-CH_4_ incubations. While recovered DNA from ^12^C incubations was considered too low for metagenome sequencing, with DNA concentrations below the limit of detection in some fractions, PCR amplification enabled characterisation of 16S rRNA gene-based community structures and comparison to the equivalent fractions from ^13^C incubations.

Six taxonomic families were represented by contigs to which a minimum of ≥1% of reads were mapped from metagenome analysis of ^13^C-enriched DNA in at least one soil. These families represented 39.8% and 83.7% of 16S rRNA gene ASVs in pH 4.5 and 7.5 samples, respectively, compared to 6.9 and 18.3% in the equivalent DNA fractions from ^12^C-incubations (Supplementary Fig. 2). Annotated metagenome and 16S rRNA amplicon libraries generated from the same ^13^C-enriched DNA contained representatives of the same C1-utilising groups, although substantial differences were observed in relative abundance. For example, while the *Methylococcaceae* was the dominant family in pH 7.5 ^13^C-enriched DNA using both approaches, it represented 5.3% (±0.3% s.e.) and 61.1% (±3.2% s.e.) of annotated contigs and 16S rRNA gene ASVs, respectively. In ^13^C-enriched pH 4.5 DNA, the dominant family in the metagenomic libraries was the *Methylocystaceae*, representing 13.9% (±1.6% s.e.) of annotated contigs but only 0.02% (±0.006% s.e.) in the amplicon libraries, where the non-methanotrophic methylotrophic *Hyphomicrobiaceae* was the most abundant at 22.0% (±5.7% s.e.). Observed differences were therefore likely due to a combination of variation in genome size and 16S rRNA gene copy number between strains in addition to the range of biases associated with different marker-gene and metagenomic sequencing approaches^2^. Nevertheless, the overall community composition determined by 16S rRNA gene amplicon libraries were highly reproducible and clearly distinct between ^12^C and ^13^C incubations, confirming that communities analysed in metagenomic libraries were enriched in methane-derived ^13^C (Supplementary Fig. 2c).

#### Predicted C1 metabolism in metagenome assembled genomes

Aerobic methylotrophic organisms utilise C1 compounds such as methane or methanol for both carbon and energy requirements^3^. Methanotrophs are one group of methylotrophs that oxidise methane to formaldehyde via methanol, which is either assimilated for generating biomass or oxidised through to CO_2_ to obtain energy and reductant. Non-methane oxidising methylotrophs, lacking methane monooxygenase, utilise methanol produced from other sources including that excreted from methanotrophs. Methane can therefore be directly and indirectly utilised by methylotrophic populations in natural communities.

To determine whether MAGs represented methanotrophic or non-methanotrophic methylotrophs, the presence of genes encoding methane monooxygenase (MMO), methanol dehydrogenases (MDH) and formate dehydrogenases (FDH) was determined after predicted protein sequence annotation. Taxonomic assignment and methylotrophic characterisation were consistent with known traits of *Methylobacter, Methylocapsa, Methylosinus* and *Methylocystis* strains, all of which possessed particulate methane monooxygenase (pMMO) and with the three *Methylosinus* MAGs also possessing soluble methane monooxygenase (sMMO). Eight MAGS lacked genes encoding an MMO but possessed MDH and FDH confirming methylotrophic capability. This included representatives of previously recognised non-methanotrophic methylotrophs including *Gemmatimonadales, Hyphomicrobium, Methylophilaceae, Methyloceanibacter* plus MAGs representative of the genera *Rudea* and *Herminiimonas*. One MAG belonged to the class *Kiritimatiellae* of the Verrucomicrobiota. While this phylum contains known methylotrophs, no pathways for C1 metabolism were identified, potentially due to the low (51.5%) estimated completeness.

### Comparison of VirSorter and DeepVirFinder in predicting virus-associated metagenome contigs

Using assembled contigs >10 kbp, metagenomic viral contigs (mVCs) were predicted using two established tools. VirSorter^4^ uses a database of viral genes plus analysis of virus-like motifs, and DeepVirFinder^5^ uses a *k-*mer based alignment-free approach, using viral genomes to train the prediction model. Both approaches provide different levels of confidence. VirSorter categories 1, 2 and 3 represent ‘most confident’, ‘likely’ and ‘possible’ virus predictions, respectively, with categories 4, 5 and 6 the equivalent for proviruses. Based on probability values, DeepVirFinder virus predictions can be considered likely (≥ 0.9, *p*-value <0.05) and probable (≥ 0.7, *p*-value <0.05)^6^. We considered the matching of CRISPR array spacers with virus protospacers as the most confident method of confirming a viral origin for an individual contig and facilitated comparison of the success of the two prediction tools. Of the 21 mVCs linked using CRISPR spacer analysis, 19 were predicted using VirSorter only, 1 was predicted using DeepVirFinder only and 1 predicted using both, the latter two being <10 kbp and identified after further analysis of 5-10 kbp contigs. Of the 270 contigs predicted as category 1, 2 or 3 mVCs by VirSorter, 49 were also predicted by DeepVirFinder, which uniquely identified a further 41 (of which only 4 were ‘likely’ viruses). These results indicate that substantially more soil virus genomes are required for training datasets using an alignment-free approach.

### Evaluation of k-mer analysis for identifying host-virus linkages

The matching of protospacers in mVCs to spacers in CRISPR arrays with 100% identity was also used to validate criteria for linkages via a ‘best hit’ homologue approach, specifically the sharing of a minimum of five homologues to the same taxonomic family. These two approaches were then compared to linkages predicted using a *k*-mer based analysis with the tool WIsH^7^, which involves the comparison and prediction of linkage of previously defined host- or virus-derived contigs on the basis of *k*-mer frequency analysis. Seven of the 21 mVCs predicted to hosts via CRISPR spacer analysis were linked to a host using WIsH and a probability score ≤ 0.05 (Supplementary Table 9), all of which were consistent at the family level. Of the 103 mVCs linked in vConTACT 2.0 analysis with an assigned host, 35 had a host predicted using WIsH, of which only 23 (66%) and 8 (23%) was the same as the homologue-based prediction at the order and family levels, respectively. These analyses therefore indicated that while the *k-*mer based approach using WIsH was partially successful in identifying correct linkages at the family level, it was not robust for identifying linkages with a high level of taxonomic resolution or confidence.

### Gene-sharing networks of viral metagenomic contigs

Predicted hosts of mVCs from this study (using CRISPR and homologue-based approaches) were generally inconsistent with the known hosts of RefSeq viruses that were linked through vConTACT 2.0 gene-sharing network analysis^8^. However, mVCs from this study placed into individual clusters all had the same predicted host (Supplementary Figure 4). Initial networks used mVCs with a conservative host prediction only (i.e. a minimum of ≥5 homologues linked to one taxonomic family). With the exception of one cluster of 3 mVCs, which were predicted to be linked to *Methylococcaceae* and non-methane oxidising methylotrophic *Hyphomicrobiaceae*, all individual networks were restricted to one family for methanotrophs or one cluster of exclusively rhizobia-linked mVCs. Further analyses included all mVCs >10 kbp in this study (i.e. including those with <5 host associated homologues) and the same networks were identified i.e. mVCs linked within the same cluster all had homologues (i.e. with <5 or ≥5) linking them to the same host family.

For *Methylocystaceae* mVCs, two separate but linked clusters were identified. The first contained CRISPR-linked *Methylocystaceae*-associated mVCs, which were recovered from both pH 4.5 and 7.5 soil (although individual mVCs were restricted to one soil pH). A second cluster was dominated by mVCs from pH 4.5 soil only, indicating that most active *Methylocystaceae* viruses belonged to one of two distinct lineages. There was a clear difference in the prediction of category-2 (‘likely’) and category-3 (‘possible’) mVCs associated with these two clusters. While category-3 viruses are often excluded prior to analysis of soil viromes^9^, CRISPR analysis demonstrated that one-third of linked mVCs were of the lowest category of confidence. It must be recognised that a proportion of the predicted category-3 mVCs in this study will not be derived from viruses, and clusters composed exclusively of category-3 mVCs without further validation (e.g. without CRISPR spacer linkages) must be interpreted with caution. However, all major clusters identified through gene-sharing network analysis contained a mixture of category-2 and −3 mVCs indicating that they represented groupings of genuine virus-derived genomes.

An intriguing finding was the linkage of host family-specific viruses from two different geographical regions (Scotland and Sweden) and contrasting soil types (agricultural loamy-sand and permafrost peatland soils), from this study and that of Emerson et al.^9^, respectively. In the latter study, 13 of 1,907 mVCs were predicted to have a methanotroph host, of which 9 were linked specifically to the *Methyocystaceae*. Intriguingly, 7 of these were also linked to a predicted *Methyocystaceae* mVC in our study, with one additional mVC predicted to have a *Methylocapsa* host (i.e. belonging to the *Beijerinckiaceae* which is another methylotrophic family of the Rhizobiales) but also contained one predicted gene with a ‘best hit’ match to a *Methylocystis* genome homologue. As soil pH is recognised as one of the dominant factors driving microbial community structures in soil^10^, it is interesting to note that linked mVCs from both studies were also from acidic soils only.

